# A least microenvironmental uncertainty principle (LEUP) as a generative model of collective cell migration mechanisms

**DOI:** 10.1101/404889

**Authors:** Arnab Barua, Josue M. Nava-Sedeño, Haralampos Hatzikirou

## Abstract

Collective migration is commonly observed in groups of migrating cells, in the form of swarms or aggregates. Mechanistic models have proven very useful in understanding collective cell migration. Such models, either explicitly consider the forces involved in the interaction and movement of individuals or phenomenologically define rules which mimic the observed behavior of cells. However, mechanisms leading to collective migration are varied and specific to the type of cells involved. Additionally, the precise and complete dynamics of many important chemomechanical factors influencing cell movement, from signalling pathways to substrate sensing, are typically either too complex or largely unknown. The question is how to make quantitative/qualitative predictions of collective behavior without exact mechanistic knowledge. Here we propose the least microenvironmental uncertainty principle (LEUP) that serves as a generative model of collective migration without incorporation of full mechanistic details. Interestingly we show that the famous Vicsek model is a special case of LEUP. Finally, as a proof of concept, we apply the LEUP to quantitatively study ofthe collective behavior of spherical *Serratia marcescens* bacteria, where the underlying migration mechanisms remain elusive.

## 1 Introduction

Collective movement of dense populations is observed in several biological systems at different scales, from massive migration of mammals ([4]) to cells during embryogenesis ([34]). In these systems, individuals which are able to propel themselves independently and interact with other nearby, start moving in a coordinated fashion once enough similar individuals are brought together. Due to the relevance of many of these processes to human activity, as well as their pervasiveness, there is a need for quantitative understanding of collective migration. *Mechanistic* models, in particular, incorporate the driving interactions between individuals in the specific system modeled. It is clear that different types of individuals, especially across different spatial scales, synchronize their movements through different mechanisms. This results in a variety of models specific to a certain individual species ([4], [17], [14]). In the specific case of biological cells, cellular migration involves numerous biophysical processes such as actin polymerization, receptor recruitment, or in bacteria, flagellar motor reversal mechanisms, to name a few [35]. However, in many cases the exact knowledge of all participating biophysical/chemical mechanisms related to a particular collective migration pattern is not trivial. Therefore, there is the need to construct models of collective migration which do not require knowledge of the exact interactions between individuals.

To address this situation, several mathematical models introduce a *phenomenological* short-range bias every individual feels. In one of the most influential collective migration models ([37]), the direction of movement of particles changes towards the mean velocity of individuals in a local neighborhood, inducing long-range swarming at the population level. Such models can be further refined into mechanistic models, where individual particle dynamics are dictated by a system of Langevin equations. In Langevin equation models, the reorientation of individual particle velocities is brought about by the existence of a local interaction potential, which is determined by neighboring particle properties. Collective migration has been achieved, for example, through the introduction of a ferromagnetic-like interaction potential, which locally aligns particle velocities polarly, or a liquid-crystal-like interaction potential, which aligns particle velocities nematically([29]).

Often neither biophysical nor phenomenological models are able to provide a plausible explanation or quantitative reproduction of collective migration patterns. Such an example is the spatiotemporal dynamics of spherical *S. marcescens* bacteria. Interestingly, prior modeling works ([2]) were able to partially reproduce the experimental results, since the underlying biophysical mechanisms are still unclear. In such cases, one could rely of machine/statistical learning methods that circumvent the biophysical details ([26];[39];[27]). However, such methods are typically of high accuracy but low interpretability, i.e. they are “black boxes” that do not offer mechanistic insights, and prone to overfitting.

From statistical mechanics, it is well-known that the precise microscopic information of a system is not needed for a macroscopic understanding of its behavior. Furthermore, entropy is the driving force behind numerous complex thermodynamic phenomena. While biological systems are subject to entropy maximization as is all matter, it is well known that living systems invest energy in decreasing their local entropy as well ([11];[38];[9]). It is, therefore, reasonable to assume that maximization as well as minimization of entropy is the driving force behind collective phenomena including migration. Under this assumption, only understanding which variables are subject to entropy optimization is required. In light of the universality of entropy optimization, we propose a potential principle that underlies the migration dynamics (see Fig. 1). Here, we postulate the Least Microenvironmental Uncertainty Principle (LEUP) for migration that aspires (i) to provide a low-dimensional statistical mechanics description, (ii) circumvent the uncertainty about the underlying biophysical mechanisms and (iii) provide a relationship to phenomenological models (e.g. the Vicsek model).

**Figure 1:**
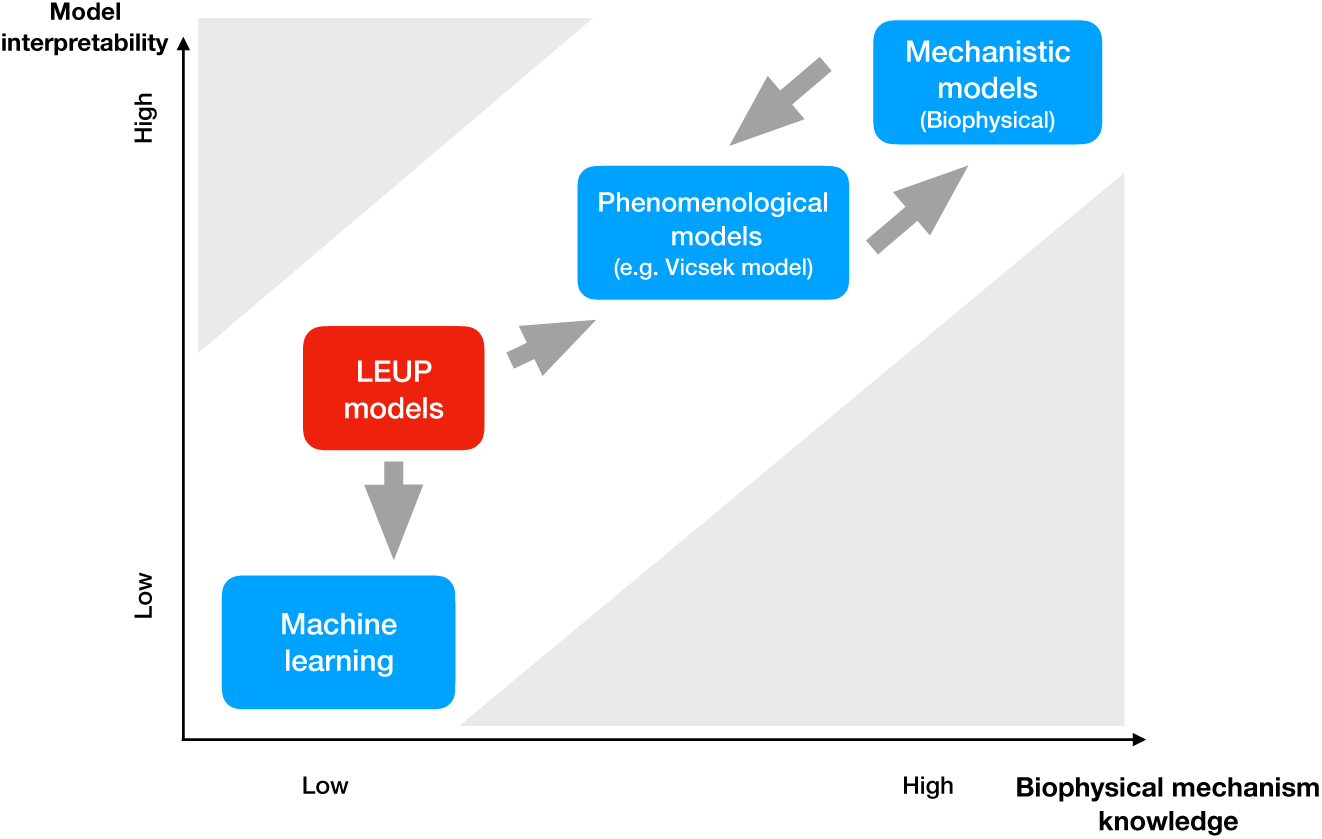
In the figure, we describe the the applicability of model classes according to amount of model interpretability and the knowledge of biophysical migration mechanisms. In the case of good knowledge of biophysical mechanisms, mechanistic models should be the natural choice. When the effects of cell-cell interaction on cell migration are only partially understood, phenomenological models are well suited. When very little is known about the system’s biophysical details, machine learning allows for the quantitative reproduction of experimental data. However, this has a toll in the interpretability of the resulting model and its generalization. LEUP models offer a great solution that allows for quantitative prediction under high mechanism uncertainty and still allow for data interpretation due to their connection with phenomenological models.

In LEUP, we view migration as an active decision-making process. It is generally accepted that many kinds of cells base their decisions according to their microenvironment ([21]; [22]; [1];[28]). More formally, the probability of cells to display certain behavior is dependent on the probability of assessing their local microenvironment. It has been argued that an energy-efficient manner for cells to encode their microenvironmental information into their decision process is by environmental entropy gradient-based decisions ([16]). In other words, a cell may react to its environment by either conforming to the information present in its environment, or acting against it. If the microenvironmental information includes information about the states of other cells, this cell-decision making process can be considered as the result of “social pressure” ([7]).

In this work, we present a Langevin model of swarming where individuals can sense the velocity orientations of other individuals in their surroundings. Individuals act as Bayesian inferrers and change their own orientation to optimize their prior, according to environmental orientation information. Under these assumptions, individuals reorient according to the entropy gradient of the environmental information. A parameter, named the sensitivity, controls the strength and directionality of the reorientation in relation to the local gradient. We find that the system adopts a steady, polar-ordered state for negative values of the sensitivity. Conversely, the system remains out of equilibrium, but partially nematic-ordered when the sensitivity is positive. Furthermore, we find that the qualitative behavior of the model depends on the values of the particle density, noise strength, sensitivity, and size of the interaction neighborhood. Finally, we showcase the LEUP principle by showing that our model replicates the collective behavior of spherical *S. marcescens* bacteria.

## 2 Materials and methods

### 2.1 The self-propelled particle framework

Moving and interacting cells are modeled by a two-dimensional self-propelled particle model (SPP). In this model, *N* ∈ ℕ cells move on a two-dimensional area. The *n*-th cell is characterized by its position, 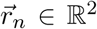, speed, *v*_*n*_ ∈ [0, *∞*) ⊂ ℝ, and an orientation *θ*_*n*_ ∈ [0, 2*π*) ⊂ ℝ. Due to the small size of cells, it is assumed that viscous forces dominate. Changes in speed and orientation result from local potentials 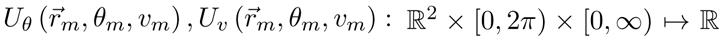 which depend on the positions and polar velocity components of cells within a radius *R ∈* ℝ_+_. The bias of the cell to follow the potential gradients is regulated by the parameters *β*_*θ*_, *β*_*v*_ ∈ ℝ, called angular and radial sensitivities, respectively. Additionally, velocity fluctuations occur due stochastic noise terms 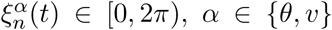 where *t ∈* ℝ_+_ denotes time. The noise will be assumed to be a zero-mean, white noise term, which has the statistical properties 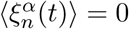 and 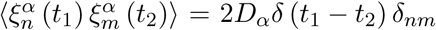, where *t*_1_ and *t*_2_ are two time points, *D*_*α*_ ∈ ℝ_+_ is either the angular (*α* = *θ*) or radial (*α* = *v*) diffusion coefficient, *δ*(*t*) is the Dirac delta, and *δ*_*nm*_ is the Kronecker delta. Finally, the radial acceleration will be assumed to be damped by a density dependent friction, *ψ*(*ρ*_*n*_). In the following, it will be assumed that the density-dependent friction is given by 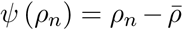, where *ρ*_*n*_ is the local cell density within the *n*-th cell’s interaction radius, and 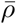 is the global average cell density. Taking everything into account, the stochastic equations of motion of the *n*-th cell read [33]

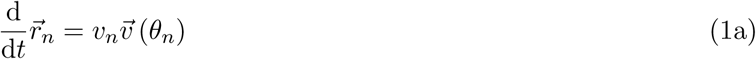

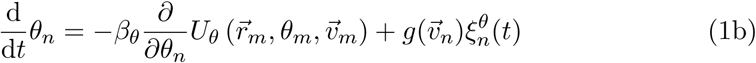

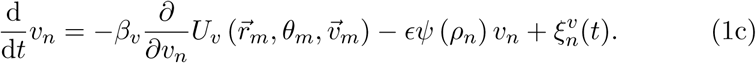

where 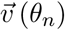 is the normalized velocity of the cell and *ϵ* is a parameter. A representation of the SPP model is shown in Fig. 2. The function 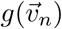 modulates the noise variance and allows us to model certain distributions (see Section 5).

**Figure 2:**
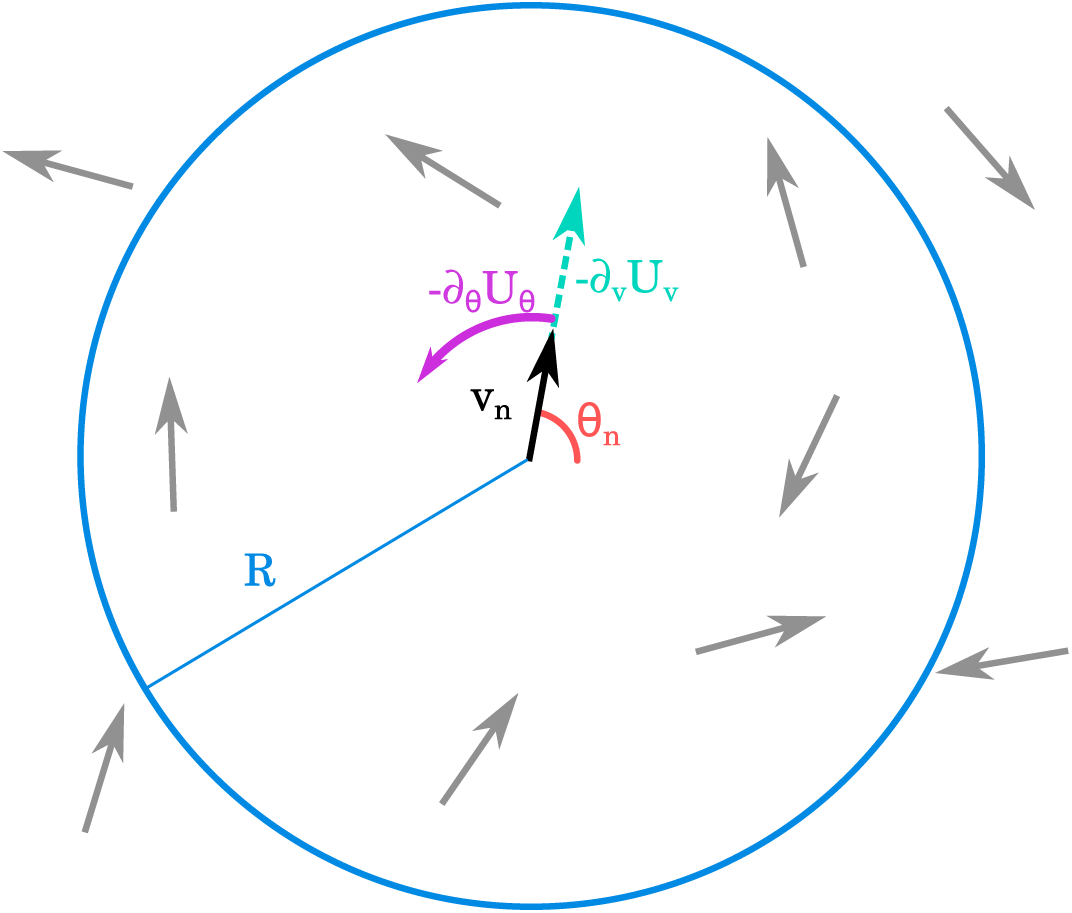
Graphic representation of the dynamics of the SPP model. The *n*-th cell is represented by a point particle with speed *v*_*n*_ and orientation *θ*_*n*_. Depending on the form of the interaction potential, the cell may feel a reorientation force −*∂*_*θ*_*U*_*θ*_ and a radial force −*∂*_*v*_*U*_*v*_ due to interaction with other cells inside the interaction neighborhood defined by the radius *R*.

The interaction potentials 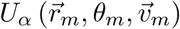, which dictate the velocity dynamics of cells, need to be specified. Biophysically, the potentials should encompass steric effects, hydrodynamic interactions, chemotactic effects, and terms arising from internal cellular processes, for example, flagellar motor dynamics, actin polymerization, receptor dynamics, etc. Finding such potentials is a formidable task since not all of the mechanisms and interactions involved are known. To circumvent this problem, a variational principle of cell decision-making related to entropy maximization [5], known as the least microenvironmental uncertainty principle (LEUP), will be used [16]. In the next section, we will discuss such a case.

### 2.2 Least microenvironmental uncertainty principle

The state of the *n*-th cell in this case is defined by its orientation *θ*_*n*_ and velocity *v*_*n*_. We assume that the orientation and velocity of cells are decoupled, i.e. one can consider orientations and velocities independent from one another. The set of intrinsic angular states of other cells within its radius of interaction is given by 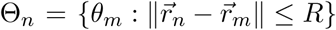, while the set of intrinsic velocity states is 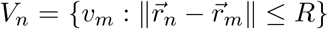. The cell reacts to the environmental information, Θ_*n*_ and *V*_*n*_, by changing its own states, *θ*_*n*_ and *v*_*n*_. The cell then acts as a Bayesian decision-maker, such that

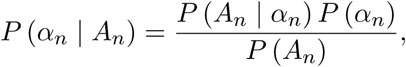

*α*_*n*_ ∈ {*θ*_*n*_, *v*_*n*_}, *A*_*n*_ ∈ {Θ_*n*_, *V*_*n*_} where *P* (*A*_*n*_ | *α*_*n*_) can be interpreted as the probability that the cell perceives all other cells in its surroundings, and *P* (*α*_*n*_) is the probability distribution of the cell’s intrinsic states (or prior). However, sensing other cells and evaluating *P* (*A*_*n*_ | *α*_*n*_) entails an energy cost. It is reasonable to assume that the cell will try to optimize its prior *P* (*α*_*n*_) for the sake of energetic frugality.

Using statistical mechanics arguments, the problem is equivalent to finding the prior that minimizes the entropy of the cell with its surroundings. Let *S* (*α*_*n*_, *A*_*n*_) be the entropy of the cell-surroundings system, *S* (*α*_*n*_) the internal entropy of the cell, and *S* (*A*_*n*_ | *α*_*n*_) the entropy of the information sensed by the cell. The entropies are connected by the relation *S* (*α*_*n*_, *A*_*n*_) = *S* (*α*_*n*_) + *S* (*A*_*n*_ | *α*_*n*_). The optimization problem is finding *P* (*α*_*n*_) that minimizes *S* (*α*_*n*_, *A*_*n*_), while making sure that *P* (*α*_*n*_) is normalized. Thus we postulate that the cells try to optimally regulate its internal states in order to minimize the uncertainty of its micro-environment:

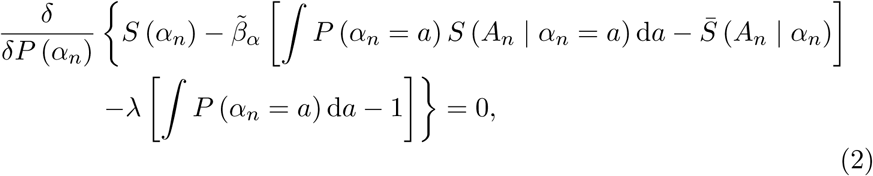

where 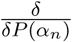 is the functional derivative, 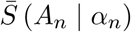 is the expected ensemble statistics, and *λ* and 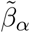 are Lagrange multipliers. Taking into account the relations among entropies, Eq. (2) yields

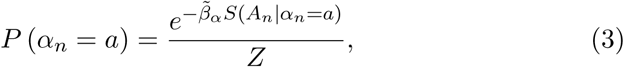

where 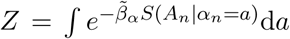, is a normalization constant, and 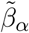 is the responsiveness of the cell. Using Eq. (3), the internal entropy of the cell, defined as *S* (*α*_*n*_) = −∫ *P* (*α*_*n*_ = *a*) ln, *P* (*α*_*n*_ = *a*) is given by

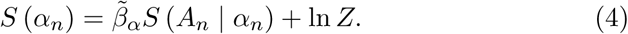

Using the relation between thermodynamic-like potentials, it is evident that the internal energy is given by

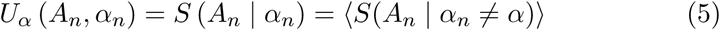

and the Helmholtz-like free energy is

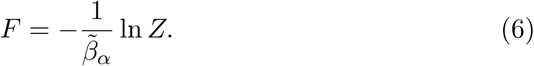

### 2.3 LEUP-based dynamics

The internal energy depends on the internal states of the cell, as well as the internal states of other cells in the surroundings. Thus, it plays the role of the required interaction potential in the equations of motion. In this case, the potential depends mainly on the cells intrinsic states. By doing so, it is evident that the responsiveness of the cells to LEUP, and the sensitivity in the equations of motion are the same. But we have chosen *β*_*α*_ as a negative sign of 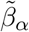 i.e. 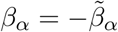. By substituting Eq. (5) into Eq. (1c), one obtains the equations of motion of the model

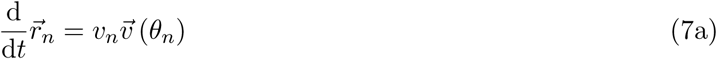

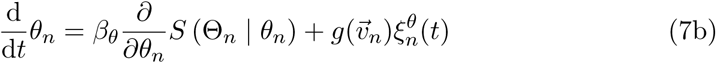

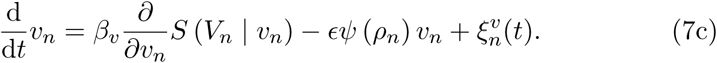

To illustrate entropy calculation, it will be assumed that the orientations of cells within the interaction neighborhood are distributed according to

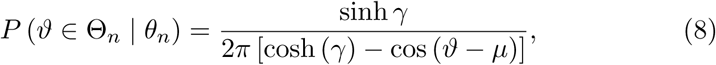

where *µ* is the mean of the distribution and *γ* is a parameter related to the variance. This is a wrapped Cauchy distribution, periodic over the interval [0, 2*π*]. Similarly, cell velocities will be assumed to be distributed half-normally

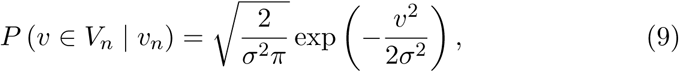

where *σ*^2^ is proportional to the variance of the distribution. Accordingly, the angular entropy is

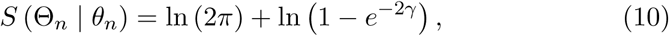

while the velocity entropy is

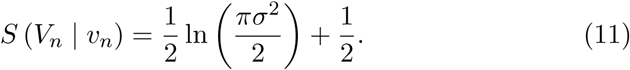

The parameter *σ* can be determined from the local speed variance, while the parameter *γ* depends on the local polar order (i.e. the degree of parallel alignment) of cell velocities in the neighborhood. In t should be noted that the qualitative behavior of the model is independent of the particular choice of distributions, and the distributions considered here are suggested only for ease of calculation. Before defining *γ*, we will first define the observables characterizing the order of the velocity field.

### 2.4 Collective migration observables

Let us define the normalized complex velocity of the *n*-th cell, *z*_*n*_ ∈ ℂ as 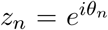, where *i* is the imaginary unit. The *k*-th moment of the velocity over an area *A* is given by 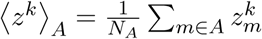, where the sum is over all cells in area *A*, and *N*_*A*_ is the total number of cells in *A*. The polar order parameter in the area *A* is given by

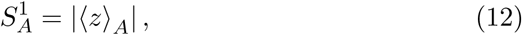

which is the modulus of the first moment of the complex velocity in *A*, while the nematic order parameter in the area *A* is given by

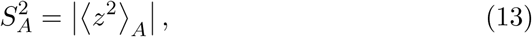

which is the modulus of the second moment of the complex velocity in *A*. The order parameters are bounded, i.e.

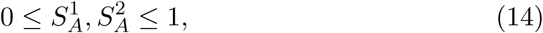

due to the complex velocities *z*_*n*_ being normalized. The parameter *γ* for the distribution of orientations in the neighborhood of the *n*-th cell is given by

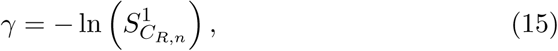

where the subindices *C*_*R,n*_ indicate a circular area of radius *R* centered at 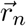.

While global polar and/or nematic order are characteristic of steady flows, rotating flow fields are commonly observed in out-of-equilibrium systems. The vorticity is an observable which is equal to twice the local angular velocity, and is thus a measure of the local strength and direction of rotation of the field. The vorticity *ω* is defined as

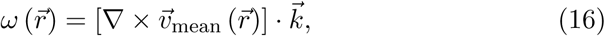

where 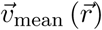 is the mean velocity field at point 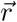, and 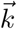 is vector normal to the plane where cells move.

### 2.5 Statistical evaluation of experiments and model predictions

For statistical evaluation, we have estimated the mean Kullbeck-Leibler divergence between experiment and simulations. We have also considered that each points in the graph from experiments and simulations are following normal distributions.

Defining the mean and the variance of each experimental data point as 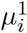 and 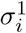, respectively, and by 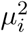 and 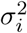 the mean and the variance of each simulated data point, then the mean K.L. divergence will be

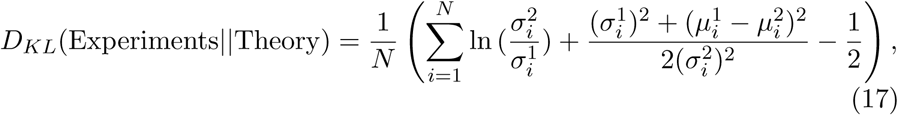

where *N* is the total number of experimental data points. We use this definition to evaluate the mean K.L. divergence in Fig. 5a and Fig. 5b. We didn’t calculate mean K.L. divergence for Fig. 5c because it’s a projection of two images i.e Fig. 5a and Fig. 5b.

**Figure 3:**
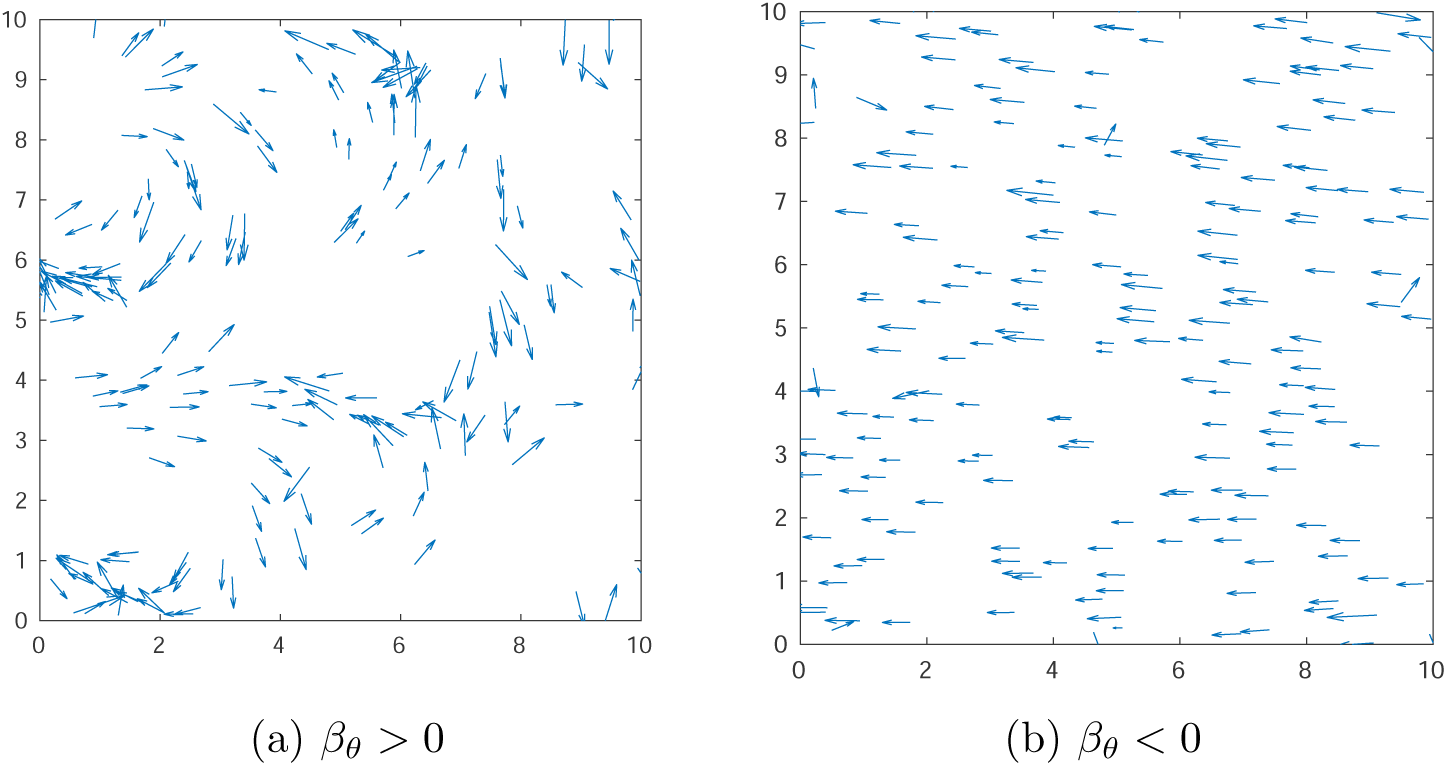
Simulation snapshots of the velocity field at long times. Arrows show the direction and magnitude of the velocity field. The snapshots were taken after 200 time steps. 200 particles were simulated, with an interaction radius of 4, and noise standard deviation of angles and speeds equal to 0.01. Here *g* = 1, *β*_*v*_ = −10 and *ϵ* = 0. In (a) the value of the angular sensitivity was equal to 25 while in (b) the angular sensitivity was equal to −5. Periodic boundaries were employed.

**Figure 4:**
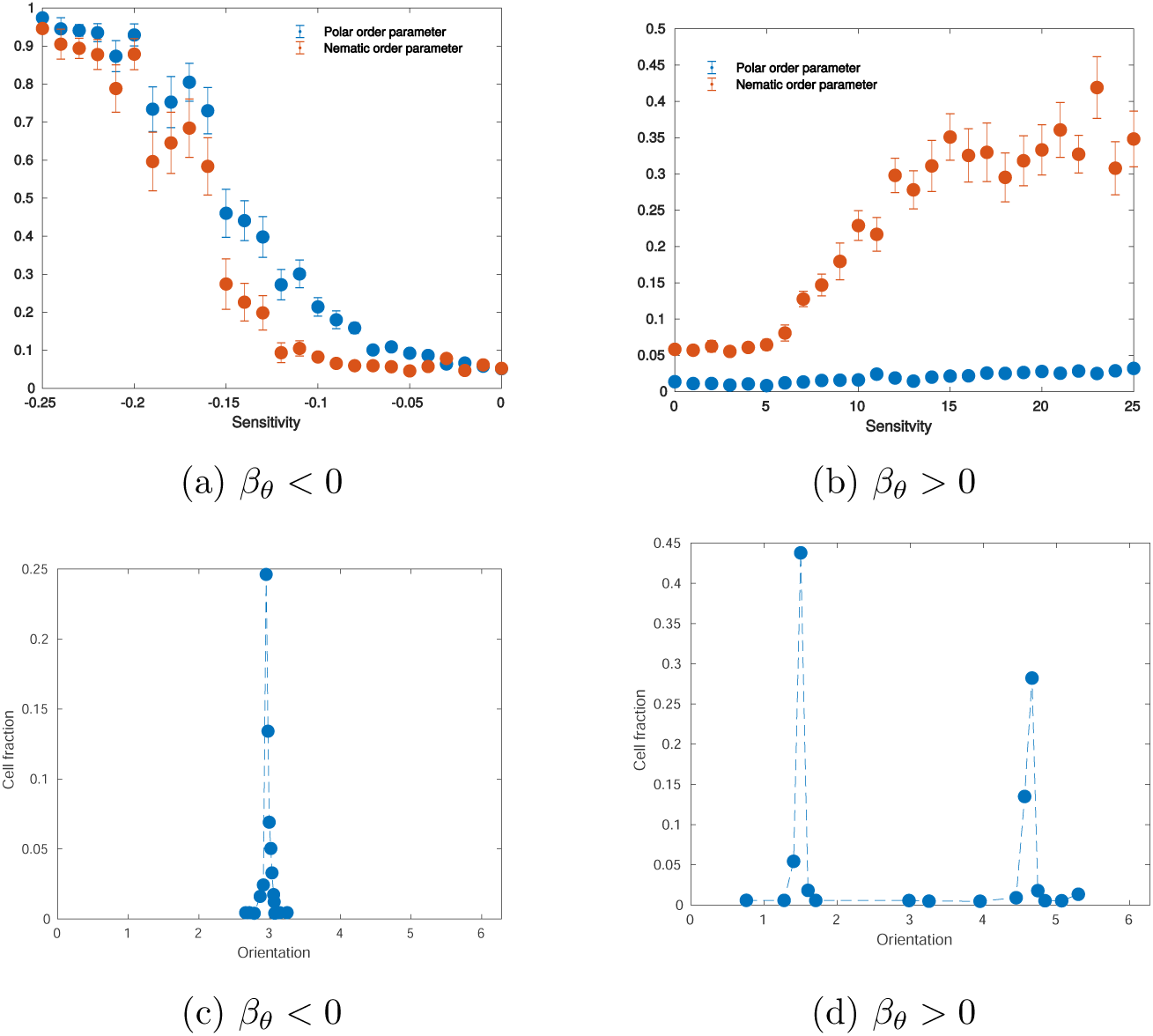
Order-disorder phase transitions and orientation distributions in two parameter regimes. Here *g* = 1, *β*_*v*_ = 0, *ϵ* = 0 and 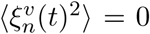. (a) In the regime *β*_*θ*_ < 0, a phase transition towards polar order occurs at a critical value of the sensitivity. (c) After the phase transition, polar order arises, and all cells have roughly the same orientation. (b) In the regime *β*_*θ*_ > 0, the phase transition towards nematic order occurs at critical value of the sensitivity. (d) There is partial nematic order after the phase transition. Accordingly, several cells have opposite orientations. (a) and (b) The number of particles was fixed at 250, noise standard deviation at 0.01, and interaction radius at 4. Values of the order parameters were averaged over 20 realizations after 1000 time steps. (c) and (d) The number of particles was fixed at 250, noise standard deviation at 0, and interaction radius at 4. The histogram was created with data from 50 realizations after 1000 time steps.

**Figure 5:**
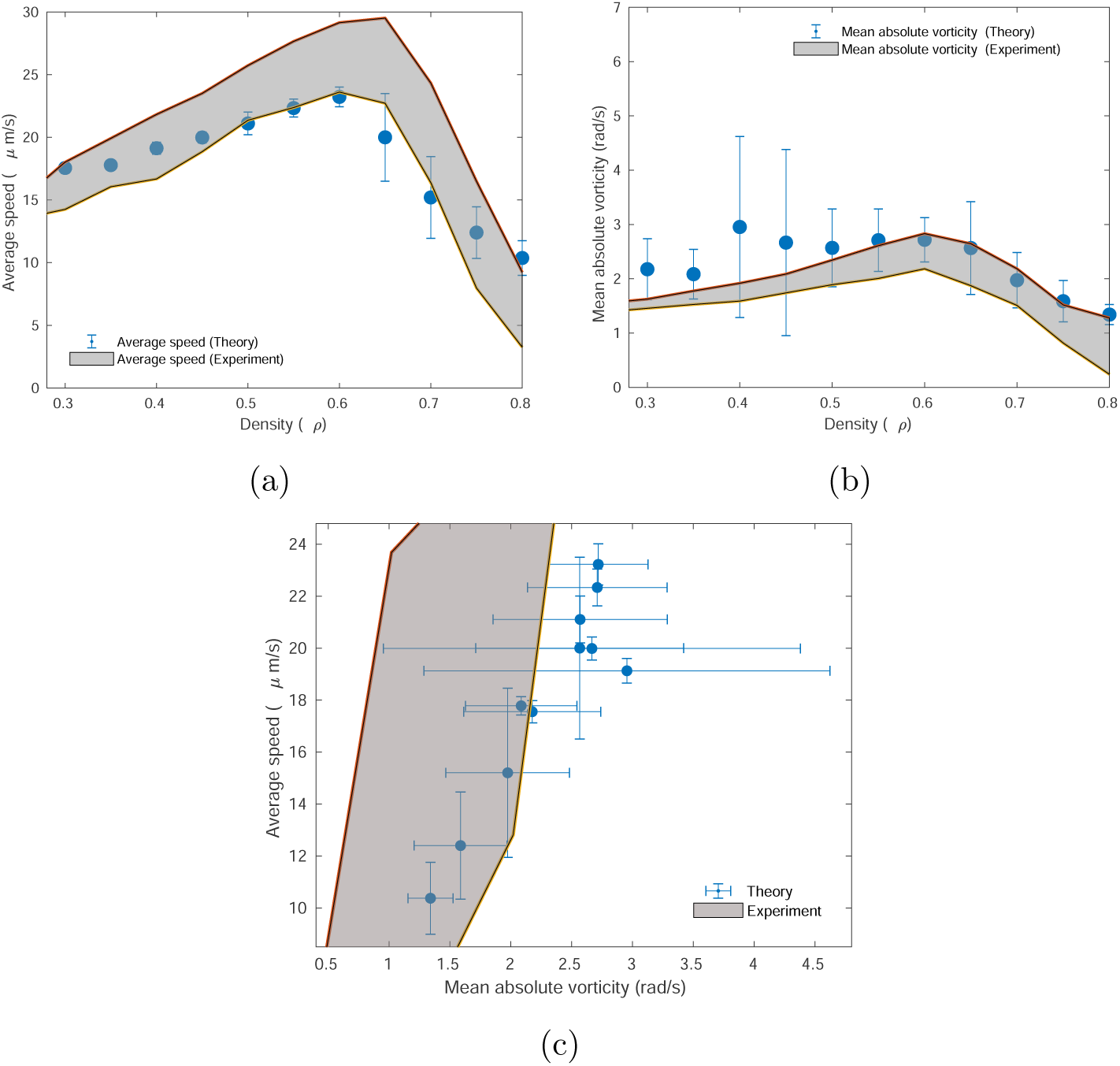
Comparison between vorticity trends in experiments and in simulations. (a) Relation between the average speed and the density. The simulation values shown are averaged over twenty realizations. (b) Dependence of the spatially normalized averaged absolute value of vorticity on the density. The simulation values shown are averaged over twenty realizations Relation between average speed versus mean absolute vorticity from simulations for various densities over twenty realizations. Experimental values were taken from [30]. Throughout all simulations, the standard deviation of the noise was set at 0.0001, interaction radius at *R* = 10, proportionality constant *ϵ* = 0.008, radial sensitivity *β*_*v*_ = −20, 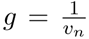 and angular sensitivity at *β*_*θ*_ = 20. Data was obtained after 500 time steps.We have also measured the Kullbeck-Leibeler divergence between experiment and theory for a) is 4.185 and for b) 1.998

## 3 Results

### 3.1 From LEUP to phenomenological models of collective migration: the relationship to the Vicsek model

By using the LEUP, we have modeled interaction as a change in velocity dictated by the local entropy gradient. The modulation of *β*_*α*_ parameters modulates the response of cells to the local entropy gradient and gives rise to relationships with know phenomenological models, such as the Vicsek model. The absolute value |*β*_*α*_| is proportional to the likelihood of the cell to change its velocity according to a given entropy gradient. If *β*_*α*_ < 0, cells tend to go against the local entropy gradient towards the entropy minimum. In the specific case of *α* = *θ*, a negative sensitivity would restrict the distribution of angles to a narrow selection. Conversely, *β*_*α*_ > 0 forces cells to follow the entropy gradient towards the entropy maximum, broadening the distribution. From here on, we will assume that the effect of cell interactions will be averaging the radial component, therefore *β*_*v*_ < 0.

To evaluate the effect of these two opposite migration strategies, we analyze the angular steady states in the two parameter regimes. Without loss of generality, we assume that, in the steady state, the mean velocity is 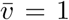. By expanding 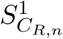 using Eq. (12), defining the components of the mean neighborhood velocity as 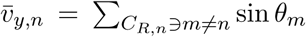 and 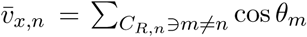, and differentiating Eq. (10), we find that the orientation of *θ*_*n*_ at the entropy extrema must be such that (see Supporting information)

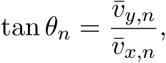

But 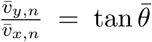, the tangent of the mean orientation of the neighbors, excluding the *n*-th cell. This results in two extremum points 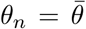 and 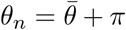, one where the velocity of the *n*-th cell is parallel to the average velocity of its neighbors, and one when it is antiparallel. In the first case

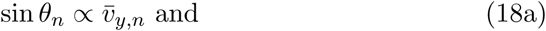

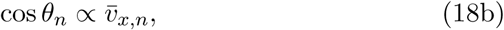

while in the second case

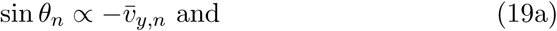

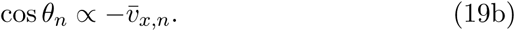

It can be shown (see Supporting information) that 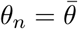 corresponds to an entropy minimum, while 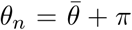 corresponds to an entropy maximum. Consequently, the behavior of **the regime** *β*_*θ*_ < 0 **is analogous to that of the Vicsek model** [37]. Conversely, the **regime** *β*_*θ*_ > 0 **corresponds to a nematic analog of the Vicsek model**.

Next, let us assume that the model has a steady state, where the Helmholtz free energy per cell is given by Eq. (6). Due to its extensivity, the Helmholtz free energy of complete, non-interacting, steady state system is

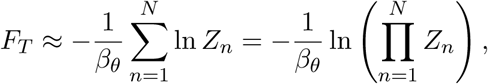

where *Z*_*n*_ is the normalization constant of Eq. (3) for the *n*-th cell. The effective normalization constant 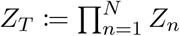 is given by

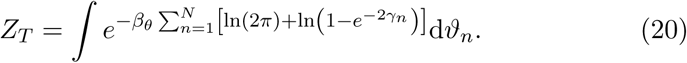

Integrating and susbtituting the resulting *Z*_*T*_ into Eq. (6) (see Supporting information), yields the Helmoltz-like free energy

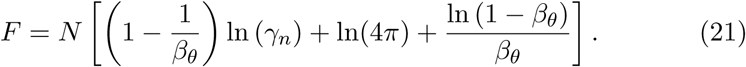

Eq. (21) is well-defined only for *β*_*θ*_ < 1. This indicates that no steady state exists for *β*_*θ*_ ≥ 1, hinting at an out-of-equilibrium regime [31]. The present model belongs to the class of models with logarithmic potentials (see Eqs. (5) and (10)). The existence of a non-normalizable state in certain parameter regimes is a staple of systems with logarithmic potentials [20].

### 3.2 Collective cell migration patterns for different parameter regimes

The model was implemented computationally to characterize the model and the effects of the different parameters on the resulting macroscopic behavior. The general qualitative behavior of the model can be observed in Fig. 3. In the regime *β*_*θ*_ < 0, cells tend to travel in a single direction after some time has elapsed, similar to the Vicsek model. Conversely, in the *β*_*θ*_ > 0 regime, cells are seen to move collectively in transient vortex-like structures, even after long times have elapsed. Qualitatively, the patterns resulting from different parameter combinations are summarized in Table 1. Analyzing simulations, two important phenomena are observed. First, there is a critical value of the interaction radius, *R*_*C*_, which separates two regimes with qualitatively different behavior. Specifically, when *β*_*θ*_ > 0, *no structures are formed for values of the interaction radius below R*_*C*_. This indicates that medium-to-long range spread of information is necessary for ordering in this regime. A second important observation is that patterns do not depend on the choice of *β*_*v*_ when this is different than zero. For *β*_*v*_ ≠ 0, LEUP dynamics divide the population into fast and slow cells. While fast cells are useful for spreading information (and therefore, increasing the effective interaction range), slow cells are necessary for maintaining local ordering. On the other hand, if we fix the initial speed distribution and assume *β*_*v*_ = 0, then we find different patterns emerging as shown in Table 1 (also see in SI).

**Table 1:**
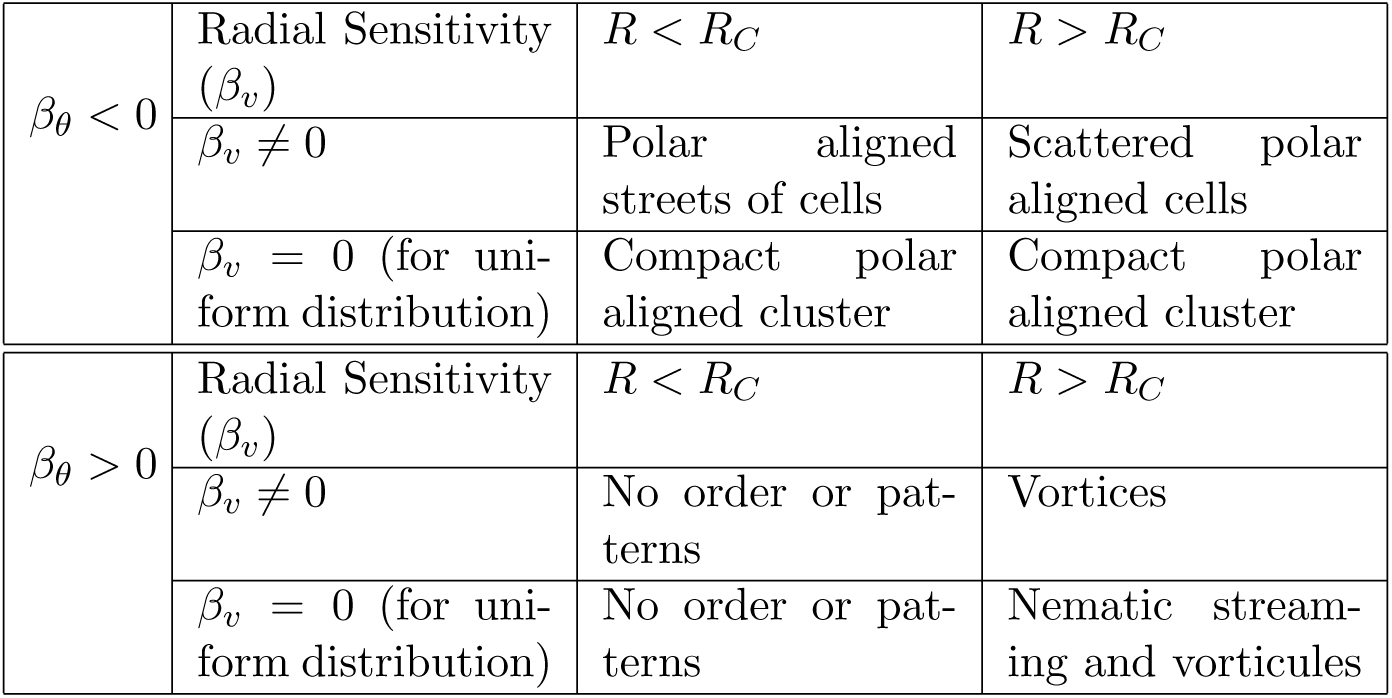
Qualitative description of the observed patterns for different angular sensitivity and interaction radius regimes, as well as radial sensitivity.

Furthermore, we quantitatively characterized global ordering at long times. The global polar order parameter, given by Eq. (12), for the complete simulation domain, measures the global degree of polar alignment, or polarization. The global nematic order parameter, given by Eq. (13) for the complete simulation domain, measures the tendency of all cells to align nematically, or along a single axis. These order parameters take a value of one when there is global order, while taking a value of zero when the system is completely disordered. It should be noted that polar order implies nematic order, but the reverse is not true.

Similarly to other velocity alignment models [29], the model shows an order-disorder transition with increasing noise amplitude and decreasing density (supplemental figure Fig. 1). More importantly, we observe that in the regime *β*_*θ*_ < 0 the system also undergoes a transition towards polar order with decreasing *β*_*θ*_. After the transition, most particles have a similar orientation (Figs. 4a and 4c). In the regime *β*_*θ*_ > 0, a phase transition is also observed towards nematic order with increasing *β*_*θ*_. In this case, however, the nematic ordering is not perfect, as evidenced by the nematic order parameter reaching values of around 0.35 after transition (Fig. 4b) compared to the value of 0.9 of the polar order parameter after transition in the *β*_*θ*_ < 0 regime. This is further evidenced by the bimodal distribution of orientations with peak separation of approximately *π* radians (Fig. 4d). These simulation results further corroborate our previous theoretical results.

In turn, we study the effect of speed sensitivity *β*_*v*_ in terms of phase transitions. We fix the angular sensitivity *β*_*θ*_, either positive or negative, thus the speed distribution will only depend on *β*_*v*_ values. When *β*_*v*_ is positive then speed distribution become bimodal and for *β*_*v*_ < 0 the speed distribution becomes unimodal (see Fig.10 in supplementary material). Moreover, we show that if the radial sensitivity *β*_*v*_ < 0 decreases then the average speed increases. On the other hand, for any value of *β*_*v*_ > 0, the first and the second moments of the speed distribution cannot be defined, since this is bimodal. Finally, for increasing cell densities, the average speed increases as well (see supplementary Fig.9).

### 3.3 A collective migration example of restricted mechanistic knowledge: the spherical bacteria case

Collective motion of bacteria has been extensively studied and modeled. Most studies have focused on the collective properties of *S. enterica, E. coli*, and *M. xanthus*. These species of bacteria are similar since they have a high aspect ratio. It has been shown that volume exclusion, coupled with a high aspect ratio, is sufficient to induce velocity alignment in the system [29], and accordingly, ordered clusters of bacteria are observed at high densities.

However, it has been recently shown [30] that even spherical *S. marcescens* bacteria do display collective migration. The biophysical mechanism whereby spherical bacteria interact with one another must be different from the high body aspect ratio volume exclusion mechanism proposed for elongated bacterial species.

Recently, a combination of biophysical agent-based and hydrodynamics model has been proposed to describe these experiments. In this study the experimental observations were only partially reproduced. Therefore, the biophysical mechanisms underlying collective migration in spherical bacteria are still not well understood. An important aspect to consider is the bacterial speed *v*_*n*_. It was found experimentally [30] that bacterial speed followed a Rayleigh distribution, dependent on bacterial density. Collective effects on cell orientations, on the other hand, were studied by observing the vortical behavior of the population [30].

To reproduce the experimentally observed Rayleigh distribution for cell speed, we chose the function 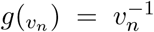 as shown in [33]. As shown in Fig. 5, our model qualitatively and quantitatively reproduces both the speed distribution and vorticity behavior of the experimental system. Interestingly, the behavior of the experimental system was replicated for high values of the sensitivities *β*_*v*_ and *β*_*θ*_, and large interaction radii *R*. Such values of the sensitivities and interaction radii indicate far-reaching, strong tendencies of bacteria to average their speeds while reorienting and traveling differently from their neighbors. Spherical, rear-propelled particles have been shown to destroy polar order as a result of hydrodynamic interactions [13], similarly to our model. Considering that *S. marcescens* is an example of a spherical, rear-propelled particle [19], our results agree with previous findings indicating that *S. marcescens* interacts through long-range hydrodynamics [36]. The long range interaction radius suggests the existence of hydrodynamically induced interaction (which has been suggested by Ariel et. al as well as by other studies ([19],[36])) or self avoiding interaction [32].

## 4 Discussion

In this work, we have introduced an off-lattice model of LEUP-induced collective migration, based on the self-propelled particles modeling framework. It was assumed that individuals changed their radial and angular velocity components independently through LEUP. Reorientation is governed by a stochastic differential equation depending on a white noise term and a force arising from an interaction potential.

The exact form of the interaction potential is very complex, and its specific form is dependent on particular details of the modeled system. While it has been shown that, in general, interactions between individuals can effectively drive the entropy of the entire system towards an extremum point [10, 15], here we do the opposite. Instead of modeling the interaction potential biophysically, it was assumed that particles followed the LEUP, which dictates that cells change their internal states in order to minimize the uncertainty of the internal states of cells in their surroundings. In this model, speeds and orientations were considered the only internal states. While cell speed was assumed to always minimize uncertainty, there was no assumption made on the cell orientation. Particles are therefore free to reorient either towards or against the gradient of entropy of the orientational distribution of particles in their neighborhood, depending on the sign of the sensitivity parameter, which also dictates the strength of the interaction. The orientational distribution in the neighborhood was assumed to be wrapped Cauchy distributed. Such a distribution facilitates the mathematical analysis of the model. However, the usage of other wrapped distributions does not qualitatively change the general behavior of the model (see Supporting information). Please note that non-parametric methods for estimating entropies without assuming any underlying parametric distributions exist. For instance, such methods employ kernel density estimation, *k*−nearest neighbours or regression methods [18].

We show that, when the parameter is negative, the model produces steady-state polar alignment patterns. Interestingly, we showed that the famous Vicsek model is a special case of LEUP. Conversely, when the parameter is positive, particles tend to reorient against the mean velocity of their neighborhood. In this regime, the free energy diverges, indicating an out-of-equilibrium parameter regime. This kind of parameter-dependent dichotomy is similarly observed in systems with logarithmic potentials [12], involved in processes such as long range-interacting gases [6], optical lattices [24], and DNA denaturation [3]. The dichotomy arises from the logarithmic form of the entropy driving interaction in our model. It has been shown that, due to the non-normalizability of the steady state solution, such systems require a time-dependent expression for their analysis [20]. Therefore, an in-depth theoretical analysis of our model would require a similar multiparticle, time-dependent expression of the angular probability densities.

As a proof of principle, we show that our model replicates the collective vortical behavior of spherical motile particles. Recently, the collective behavior of spherical particles been modeled as a combination of steric repulsion and hydrodynamic interactions [23]. Our study has shown that hydrodynamics and steric interactions induce long-range microenvironmental entropy maximization, which coincides with the *β*_*θ*_ > 0 LEUP regime. This generalizes the type of biophysical mechanisms required to produce vortical patterns.

It should be noted that, while spherical *S. marcescens* bacteria have been modeled biophysically, their collective behavior has not been completely reproduced [2]. This hints at an additional biological and/or biochemical interaction between cells. While our LEUP-based model is coarse-grained in terms of specific biophysical/biochemical interactions, it allows for successfully reproducing the experimentally observed collective velocity behavior by fitting a only few parameters. The application to spherical bacteria allows us to showcase the potential of the LEUP principle when the precise interaction mechanisms are not known.

As already mentioned, we have made some assumptions to simplify the model. Our model assumes a Gaussian, white noise term in the SDEs. This results in normal diffusive behavior in the absence of interactions. It has been observed experimentally, however, that in some conditions, cells perform Lévy walks resulting in superdiffusive behavior [25]. By changing the distribution or time correlations of the noise [8, 26], it would be possible to both replicate the non-Gaussian dynamics of single cells, and investigate the effect of single anomalous dynamics on collective behavior.

We have also assumed that particle velocities are the only internal states relevant for reorientation, for simplicity and as a proof of concept of the LEUP principle. However, it is reasonable to think that other states, such as relative position or adhesive state, may be relevant to include when modeling specific systems. It remains to be seen how additional states may impact the order and emerging patterns of the system.

As stated above, LEUP circumvents the biophysical details of cell migration and allows for replicating a plethora of collective migration patterns. For instance, we have analytically derived the polar and nematic alignment Vicsek models for LEUP arguments. In this sense, LEUP acts as a generative model for collective migration mechanisms. This is particularly useful upon limited knowledge of such mechanisms, a problem called structural model uncertainty. Another advantage of LEUP is the mapping of biophysical mechanism combination to the *β* > 0 or *β* < 0 regimes. This allows for unifying the model analysis but for a better classification of migration mechanisms. Finally, known mechanisms or data could be easily integrated to our proposed framework by further constraining the LEUP dynamics.

## Acknowledgements

AB thanks the International Graduate School of HZI, Braunschweig. JMNS thanks the Center for Information Services and High Performance Computing (ZIH) at TU Dresden for providing an excellent infrastructure. The authors would like to thank Andreas Deutsch and Rainer Klages for their helpful comments and fruitful discussions. JMNS acknowledges support from the joint scholarship program DAAD-CONACYT-Regierungsstipendien (50017046), and from ROCKET (031L0139B). H.H. would like to acknowledge the support by MicMode-I2T (01ZX1710B) and H.H. is supported by SYSIMIT (01ZX1308D) and MulticellML (01ZX1707C) by the Federal Ministry of Education and Research (BMBF) and by the SYSMIFTA (031L0085B) of the ERACOSYSMED initiative.

## Notes

#### Summary of Updates

-New Figure - Revised text and abstract

