## Supplementary material for "A least microenvironmental uncertainty principle (LEUP) as a generative model of collective cell migration mechanisms"

September 4, 2019

#### Calculation of the curvature of micro-environmental entropy and Helmholtz free energy

The entropy of the wrapped Cauchy distribution is

$$\begin{aligned}S\left(\Theta_n \mid \theta_n\right) &= \ln\left(2\pi\right) \\ &+ \ln\left\{1 - \frac{1}{N_{C_{R,n}}^2}\left[\bar{v}_{y,n}^2 + \bar{v}_{x,n}^2 + 2\left(\bar{v}_{y,n}\sin\theta_n + \bar{v}_{x,n}\cos\theta_n\right)\right]\right\}.\end{aligned} \tag{1}$$

To discriminate the maximum from the minimum point, the value of $\frac{\partial^2}{\partial\theta_n^2} S\left(\Theta_n \mid \theta_n\right)$ must be evaluated at each extremum point. The second derivative is given by

$$\begin{aligned}\frac{\partial^2}{\partial\theta_n^2} S\left(\Theta_n \mid \theta_n\right) =& \frac{\frac{2}{N_{C_{R,n}}^2}\left(\bar{v}_{y,n}\sin\theta_n + \bar{v}_{x,n}\cos\theta_n\right)}{1 - \frac{1}{N_{C_{R,n}}^2}\left[\bar{v}_{y,n}^2 + \bar{v}_{x,n}^2 + 2\left(\bar{v}_{y,n}\sin\theta_n + \bar{v}_{x,n}\cos\theta_n\right)\right]} \\ &- \left\{\frac{\frac{2}{N_{C_{R,n}}^2}\left(\bar{v}_{y,n}\cos\theta_n - \bar{v}_{x,n}\sin\theta_n\right)}{1 - \frac{1}{N_{C_{R,n}}^2}\left[\bar{v}_{y,n}^2 + \bar{v}_{x,n}^2 + 2\left(\bar{v}_{y,n}\sin\theta_n + \bar{v}_{x,n}\cos\theta_n\right)\right]}\right\}^2.\end{aligned}$$

In an extremum point, $\frac{\partial}{\partial\theta_n} S\left(\Theta_n \mid \theta_n\right) = 0$, therefore

$$\left.\frac{\partial^2}{\partial\theta_n^2} S\left(\Theta_n \mid \theta_n\right)\right|_{\text{ext}} = \frac{\frac{2}{N_{C_{R,n}}^2}\left(\bar{v}_{y,n}\sin\theta_n + \bar{v}_{x,n}\cos\theta_n\right)}{1 - \frac{1}{N_{C_{R,n}}^2}\left[\bar{v}_{y,n}^2 + \bar{v}_{x,n}^2 + 2\left(\bar{v}_{y,n}\sin\theta_n + \bar{v}_{x,n}\cos\theta_n\right)\right]}. \tag{2}$$

Defining $\kappa$ as the proportionality constant relating $\sin\theta_n$ and $\cos\theta_n$ with $\bar{v}_{y,n}$ and $\bar{v}_{x,n}$, respectively, the second derivative evaluated at $\theta_n = \bar{\theta}$ is

$$\left.\frac{\partial^2}{\partial\theta_n^2} S\left(\Theta_n \mid \theta_n\right)\right|_{\theta_n=\bar{\theta}} = \frac{\frac{2\kappa}{N_{C_{R,n}}^2}\left(\bar{v}_{y,n}^2 + \bar{v}_{x,n}^2\right)}{1 - \left(S_{C_{R,n}}^1\right)^2} > 0, \tag{3}$$

because the numerator is positive definite, and the denominator is positive given the bounds of the order parameters. The extremum point $\theta_n = \bar{\theta}$ therefore corresponds to an entropy minimum. Consequently, the behavior of the regime $\beta < 0$ is analogous to that of the Vicsek model. Conversely, at $\theta_n = \bar{\theta} + \pi$ we find that

$$\frac{\partial^2}{\partial \theta_n^2} S\left(\Theta_n \mid \theta_n\right)\Big|_{\theta_n=\bar{\theta}} = \frac{\frac{-2\kappa}{N_{C_{R,n}}^2}\left(\bar{v}_{y,n}^2 + \bar{v}_{x,n}^2\right)}{1-\left(S_{C_{R,n}}^1\right)^2} < 0, \tag{4}$$

using the same arguments as for the $\theta_n = \bar{\theta}$ point. Therefore, the point $\theta_n = \bar{\theta}+\pi$ corresponds to the entropy maximum. Then, the regime $\beta > 0$ corresponds to a nematic analog of the Vicsek model. Next, let us assume that the model has a steady state, where the Helmholtz free energy per bacterium is given by $F = -\frac{1}{\beta_\theta}\ln Z$. Due to its extensivity, the Helmholtz free energy of the complete system is

$$F_T = -\frac{1}{\beta_\theta}\sum_{n=1}^{N}\ln Z_n = -\frac{1}{\beta_\theta}\ln\left(\prod_{n=1}^{N} Z_n\right),$$

where $Z_n$ is the normalization constant of $n$-th cell( see Eq.(3) in paper ).

The effective normalization constant $Z_T \coloneqq \prod_{n=1}^{N} Z_n$ is given by

$$Z_T = \int e^{-\beta_\theta \sum_{n=1}^{N}\left[\ln(2\pi)+\ln\left(1-e^{-2\gamma_n}\right)\right]}\mathrm{d}\vartheta_n. \tag{5}$$

The integration is performed over the orientations of all cells in the system. Moreover, the dependency of each $\gamma_n$ on all angles $\theta_n$ is complex and makes integration challenging. However, variation of $\theta_n$ for all $n$ translates into a variation in all $\gamma_n$. Therefore, Eq. 5 is equivalent to

$$Z_T = \int e^{-\beta_\theta \sum_{n=1}^{N}\left[\ln(2\pi)+\ln\left(1-e^{-2\gamma_n}\right)\right]}\mathrm{d}\gamma_n. \tag{6}$$

Expanding up to linear terms around $\gamma_n = 0$ yields

$$Z_T = \int e^{-\beta_\theta \sum_{n=1}^{N}\left[\ln(2\pi)+\ln(2\gamma_n)\right]}\mathrm{d}\gamma_n,$$

which after rearranging terms and integrating reduces to

$$Z_T = \left[\frac{1}{(4\pi)^{\beta_\theta}}\frac{\gamma_n^{1-\beta_\theta}}{1-\beta_\theta}\right]^N. \tag{7}$$

Substituting Eq. 7 into the expression of the Helmholtz free energy (Eq. (6) in the main text), and rearranging terms, yields the Helmholtz free energy

$$F = N\left[\left(1-\frac{1}{\beta_\theta}\right)\ln\left(\gamma_n\right) + \ln(4\pi) + \frac{\ln\left(1-\beta_\theta\right)}{\beta_\theta}\right]. \tag{8}$$

Table 1: Qualitative description of the observed patterns for different sensitivty and interaction radius regimes, as well as speed distributions ($\beta_v = 0$)

| | | | |
|---|---|---|---|
| $\beta_\theta < 0$ | Speed distribution | $R < R_C$ | $R > R_C$ |
| | Delta distribution | Polar aligned streets of cells | Scattered polar aligned cells |
| | Uniform distribution | Compact polar aligned cluster | Compact polar aligned cluster |
| | Rayleigh distribution | Compact polar aligned cluster | Compact polar aligned cluster |
| $\beta_\theta > 0$ | Speed distribution | $R < R_C$ | $R > R_C$ |
| | Delta distribution | No order or patterns | Nematic streaming |
| | Uniform distribution | No order or patterns | Nematic streaming and vorticules |
| | Rayleigh distribution | No order or patterns | Vortices |

Eq. 8 is well-defined only for $\beta_\theta < 1$. This indicates that no steady state exists for $\beta_\theta \geq 1$, hinting at an out-of-equilibrium regime The present model belongs to the class of models with logarithmic potentials.
The existence of a non-normalizable state in certain parameter regimes is a staple of systems with logarithmic potentials.

### Pattern formation in different $\beta_\theta$ regime (see Table 1) ($\beta_v = 0$)

We have defined the range of the polar order parameter as P, nematic order parameter as N and mean absolute vorticity as V.

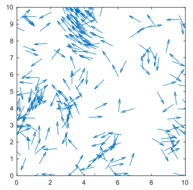

(a) Polar aligned street cells (P=0.6 to 0.7, N=0.5 to 0.7, V=0 to 0.04)

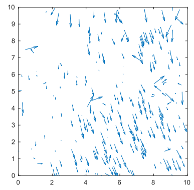

(b) Compact polar aligned cells (P=0.8 to 1.0, N=0.7 to 1.0, V=0 to 0.065)

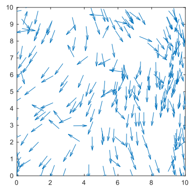

(c) Scattered polar aligned cells (P=0.7 to 1.0, N=0.7 to 1.0, V=0 to 0.05)

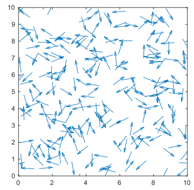

(d) No order or patterns (P=0 to 0.09, N=0 to 0.07, V=0 to 0.065)

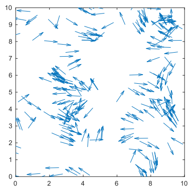

(e) Nematic streaming (P=0 to 0.05, N=0.3 to 0.5, V=0 to 0.065)

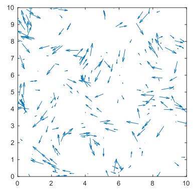

(f) Nematic streaming and vorticules (P=0 to 0.03, N=0.2 to 0.4, V=0 to 0.04)

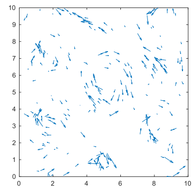

(g) Vortices (P=0 to 0.03, N=0.2 to 0.4, V=0.1 to 0.35)

Figure 1: All type of patterns are captured for different $\beta_\theta$ values. Patterns have been changed due to velocity distributions and interaction radius.

#### Polar order parameter in angular sensitivity ($\beta_\theta < 0$) regime

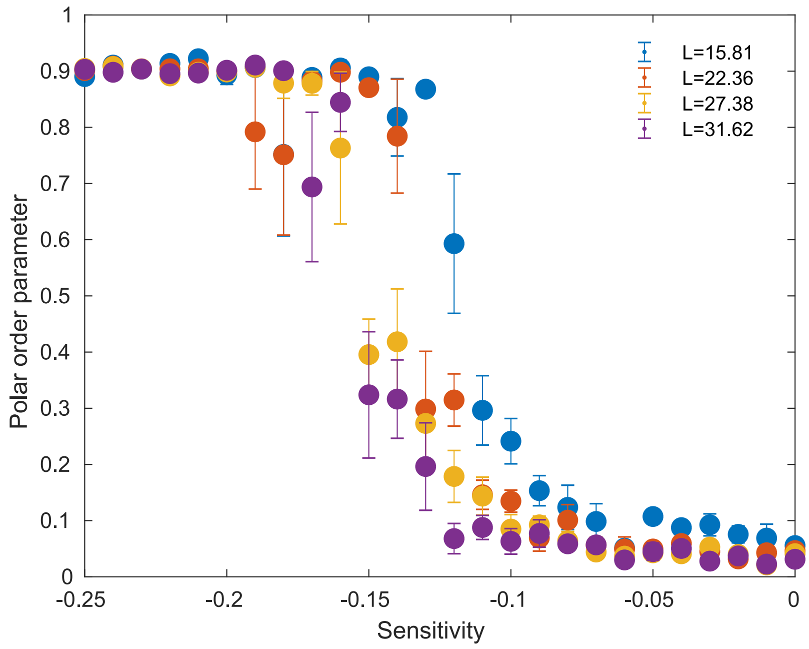


Figure 2: Polar order parameter vs. angular sensitivity graph. Where interaction radius is at 3, and standard deviation of noise is at 0.1. Density is fixed at 1.0. Here micro-environmental entropy has been taken from wrapped Cauchy distribution. Here $g = 1, \beta_v = 0$, $\epsilon = 0$ and $\langle \xi_n^v(t)^2 \rangle = 0$ . All the order parameters were averaged over 5 realizations after $10^3$ time steps.

#### Polar order parameter vs. angular noise graph

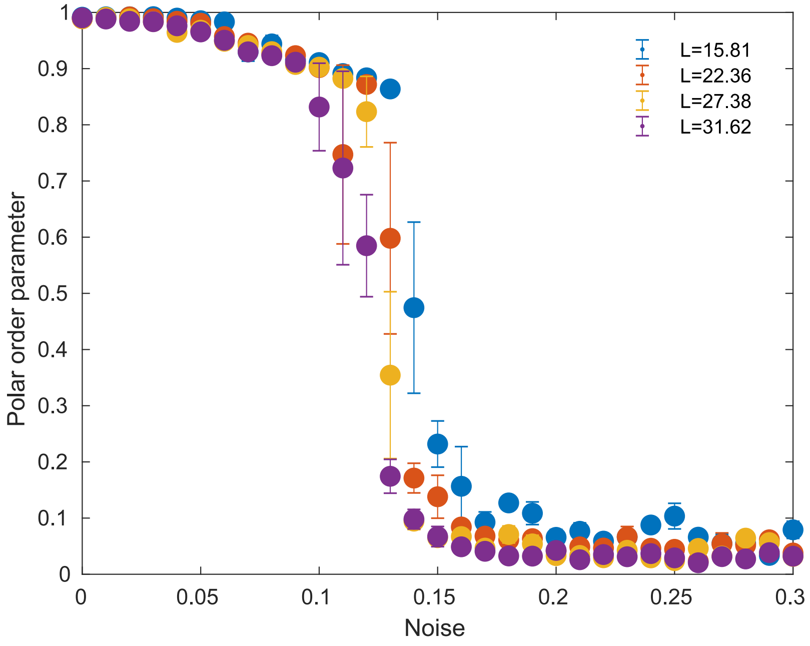


Figure 3: Polar order parameter vs. angular noise graph. Where interaction radius is at 3 and angular sensitivity is at -0.2. Density is fixed at 1.0. Here micro-environmental entropy has been taken from wrapped Cauchy distribution. Here $g = 1, \beta_v = 0$, $\epsilon = 0$ and $\langle \xi_n^v(t)^2 \rangle = 0$ . All the order parameters were averaged over 5 realizations after $10^3$ time steps.

#### Nematic order parameter in angular sensitivity ($\beta_\theta > 0$) regime

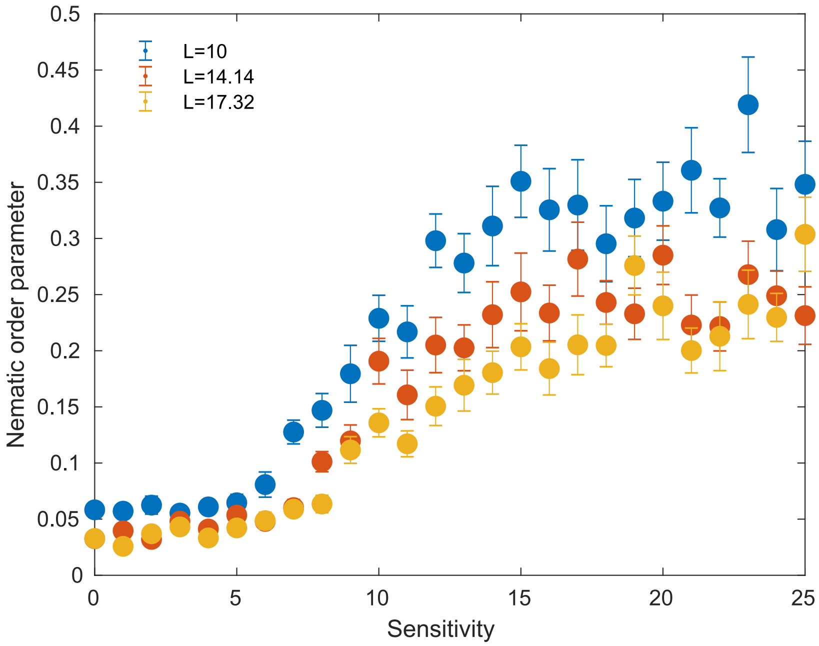


Figure 4: Nematic order parameter vs. angular sensitivity graph. Where interaction radius is at 3 and standard deviation of angular noise is at 0.05. Density is fixed at 2.5. Here micro-environmental entropy has been taken from wrapped Cauchy distribution. Here $g = 1, \beta_v = 0$, $\epsilon = 0$ and $\langle \xi_n^v(t)^2 \rangle = 0$ . All the order parameters were averaged over 20 realizations after $10^3$ time steps.

#### Nematic order parameter vs. angular noise graph

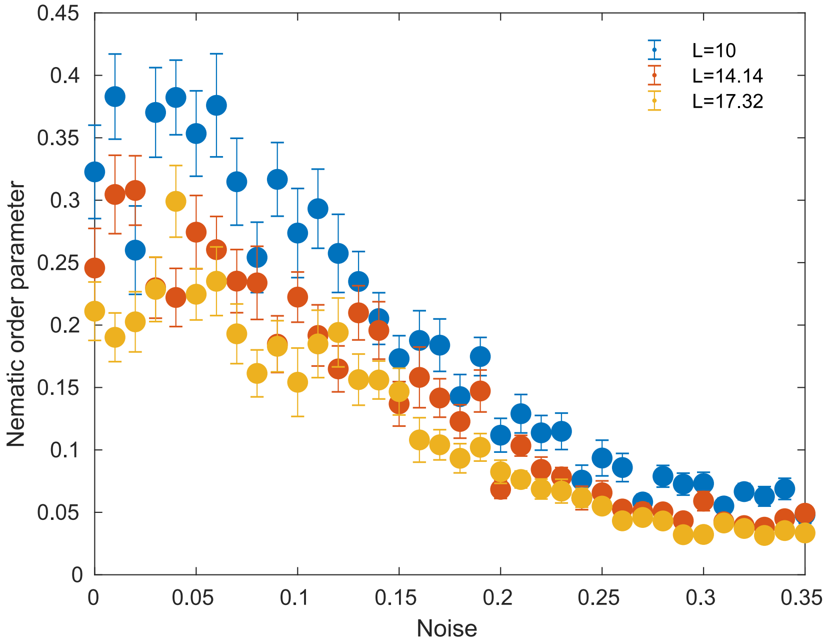


Figure 5: Nematic order parameter vs. angular noise graph. Where interaction radius is at 3 and sensitivity is at 20.Density is fixed at 2.5. Here microenvironmental entropy has been taken from wrapped Cauchy distribution. Here $g = 1, \beta_v = 0$, $\epsilon = 0$ and $\langle \xi_n^v(t)^2 \rangle = 0$ . All the order parameters were averaged over 20 realizations after $10^3$ time steps.

#### Order parameters vs. density graph

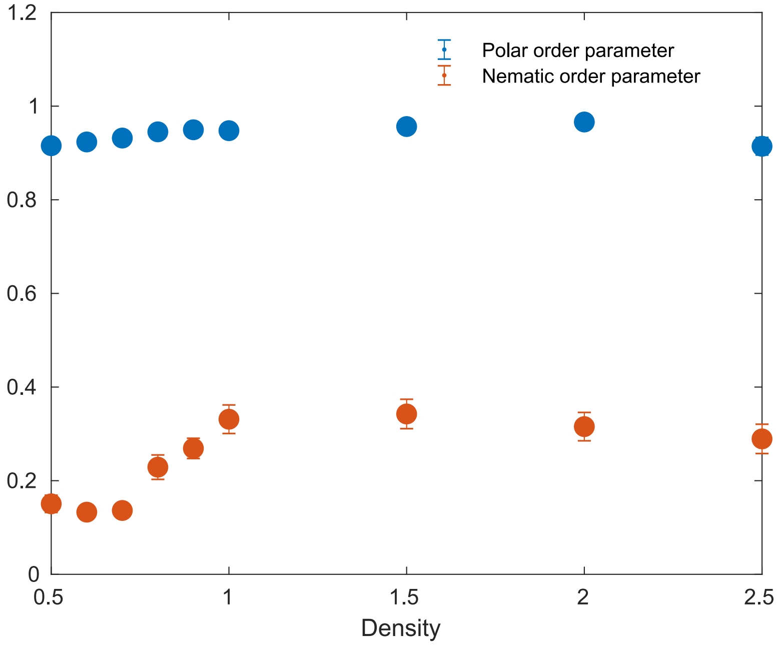


Figure 6: Polar order parameter and nematic order parameter vs. density graph. Where interaction radius is at 3 and sensitivity is fixed at 15(for nematic order parameter) and -0.2 (for polar order parameter). Here micro-environmental entropy has been taken from wrapped Cauchy distribution. Here $g = 1, \beta_v = 0$, $\epsilon = 0$ and $\langle \xi_n^v(t)^2 \rangle = 0$ . All the order parameters were averaged over 20 realizations after $10^3$ time steps.

#### Polar order parameter and nematic order parameter in angular sensitivity ($\beta_\theta < 0$) regime

When we took micro-environmental entropy from wrapped exponential distribution we didn't find any qualitative change in phase transition phenomena in $\beta_\theta < 0$ regime.

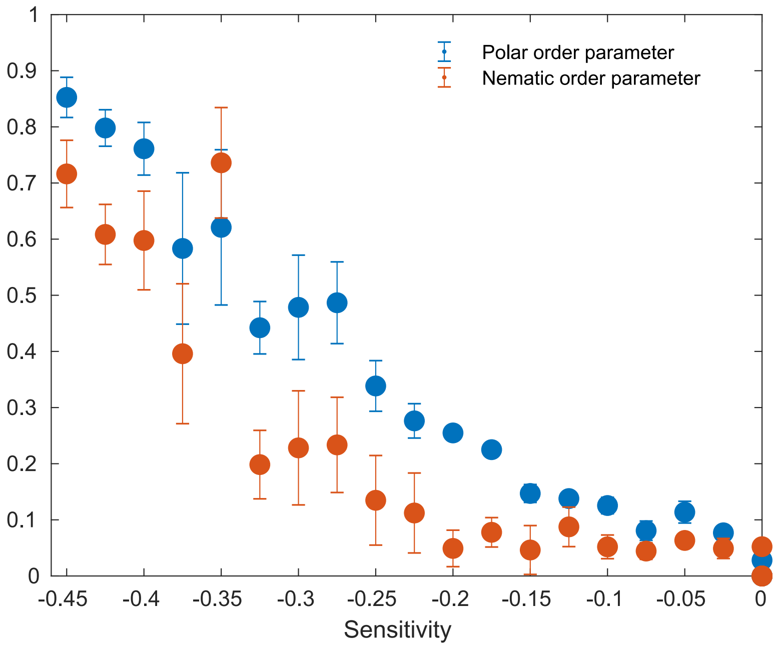


Figure 7: Polar order parameter and nematic order parameter vs. angular sensitivity ($\beta_\theta < 0$) graph. Where interaction radius is at 3, and standard deviation of noise is at 0.05. Density is fixed at 2.5. Here micro-environmental entropy has been taken from wrapped exponential distribution. Here $g = 1, \beta_v = 0$, $\epsilon = 0$ and $\langle \xi_n^v(t)^2 \rangle = 0$ . All the order parameters were averaged over 5 realizations after 250 time steps.

#### Polar order parameter and nematic order parameter in angular sensitivity ($\beta_\theta > 0$) regime

When we took micro-environmental entropy from wrapped exponential distribution, we didn't find any qualitative change in phase transition phenomena in $\beta_\theta > 0$ regime.

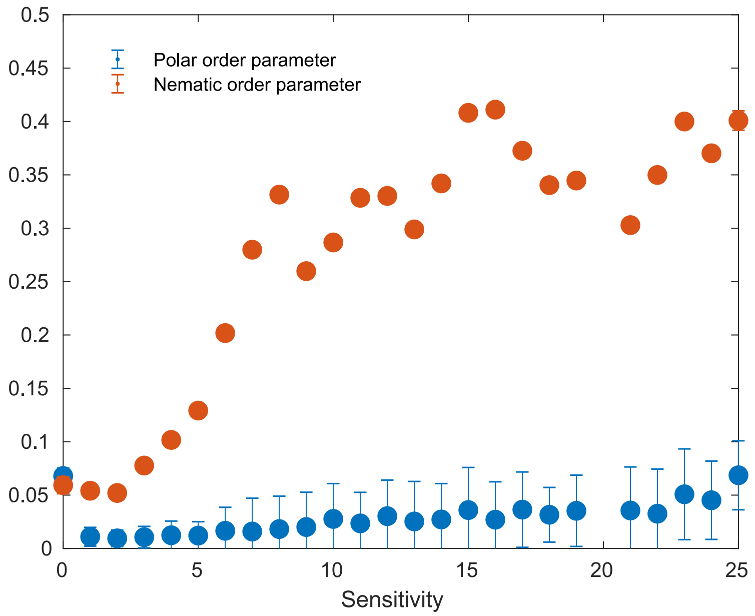


Figure 8: Polar order parameter and nematic order parameter vs. angular sensitivity ($\beta > 0$) graph. Where interaction radius is at 3, and standard deviation of noise is at 0.05. Density is fixed at 2.5. Here micro-environmental entropy has been taken from wrapped exponential distribution. Here $g = 1, \beta_v = 0$, $\epsilon = 0$ and $\langle \xi_n^v(t)^2 \rangle = 0$ . All the order parameters were averaged over 20 realizations after $10^3$ time steps.

#### Average speed vs. radial sensitivity

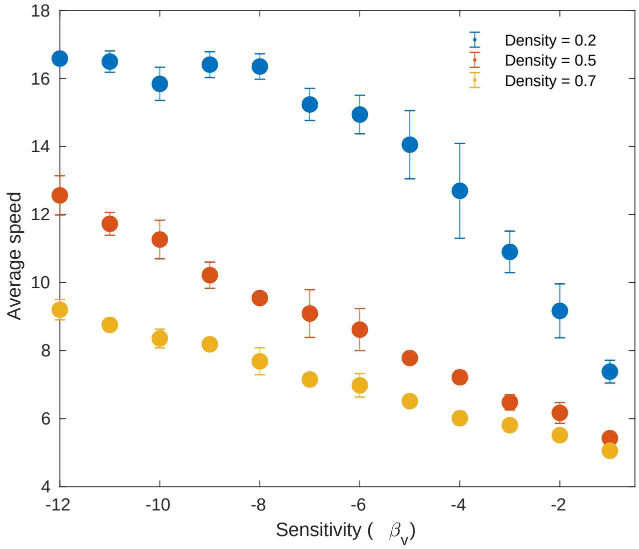


Figure 9: Average speed vs. radial sensitivity ($\beta_v$) for different densities. The standard deviation of angular noise was fixed at 0.01, $\beta_\theta = 20$, $\psi = 0$, the box size has been fixed to 30 and the interaction radius was $R = 10$.

#### Probability distributions for speed

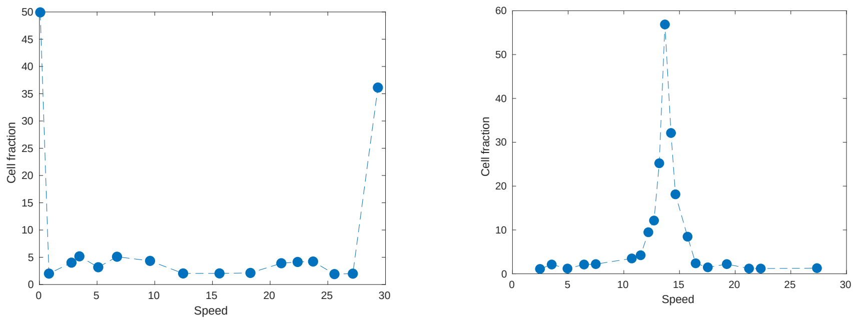


(a) $\beta_v > 0$

(b) $\beta_v < 0$

Figure 10: The number of particles was fixed at 200, noise standard deviation at 0.05, $\beta_\theta$ is at -2 and interaction radius at 4. For (a) and (b) $\beta_v$ were fixed at 25 and -10. Simulations were averaged over 15 realizations after 200 time steps.
